# Intersecting Transcriptomic Landscapes of Hypertension and Kidney Function in African American Women

**DOI:** 10.1101/2025.01.16.633488

**Authors:** Malak Abbas, Pamela Martin, Merry Lindsey, Eric S. Bennett, Thomas L. Brown, Chike Nzerue, Clintoria R. Williams, Amadou Gaye

## Abstract

**Background:** Hypertension is a major risk factor for chronic kidney disease (CKD) and disproportionately affects African American women, contributing to disparities in kidney health outcomes. The biological mechanisms connecting hypertension to reduced kidney function, particularly in understudied populations, remain poorly understood. This study leverages transcriptomic analyses to uncover shared molecular signatures associated with hypertension and kidney function, focusing on female-specific profiles.

**Methods:** The study analyzed whole-blood mRNA sequencing data from a cohort of 344 African American women, divided equally into discovery (n = 172) and validation (n = 172) datasets, along with 147 African American men. Differential expression (DE) analyses were performed to identify mRNAs associated with hypertension and kidney function (measured as eGFR). Female-specific findings were determined by comparing results between females and males. Pathway enrichment analyses were subsequently conducted to link the identified mRNAs to key biological mechanisms.

**Results:** Comparative analyses revealed unique transcriptomic profiles in females, underscoring the role of sex-specific factors in disease progression. DE analyses identified 95 female-specific genes associated with both hypertension and eGFR. Subsequent pathway enrichment analysis with the 95 genes revealed key pathways related to fibrosis, inflammation, lipid metabolism, and endothelial dysfunction. The list of 95 includes TGF-β1 and PNPLA2 implicated in fibrotic and metabolic dysregulation, and immune system players such as IL32 and TNFSF12 that amplify inflammation and kidney injury.

**Conclusions:** This study provides novel insights into the transcriptomic mechanisms underlying hypertension and kidney function in African American women. The findings emphasize the importance of addressing sex-specific and population-specific molecular mechanisms to inform precision medicine approaches and reduce health disparities hypertension-related impaired kidney function. Future research should prioritize experimental validation and longitudinal studies to further elucidate these pathways.

## INTRODUCTION

Hypertension is a leading risk factor for chronic kidney disease (CKD), cardiovascular complications, and premature mortality globally ^1^. The World Health Organization estimated that 1.28 billion adults between the ages of 30 and 79 globally are affected by hypertension, with approximately two-thirds residing in low- and middle-income countries ^2^.

In the United States, approximately one-third of adults, around 75 million people, are affected by hypertension ^3^. Of these, nearly half have hypertension that remains uncontrolled ^4^. Hypertension is more prevalent among African Americans compared to other racial and ethnic groups in the United States ^5^. Data from the National Health and Nutrition Examination Survey (NHANES) indicate that the age-adjusted prevalence of hypertension is approximately 54% in African American men and 57% in African American women, while the corresponding rates for white men and women are 46% and 48%, respectively ^6,7^.

Among African American women, the burden of hypertension is disproportionately higher ^6^, contributing to significant disparities in kidney health outcomes, including the accelerated decline in glomerular filtration rate and higher rates of end-stage renal disease ^8,9^. Despite the clinical importance of this population, research on the genomics of hypertension and its intersection with kidney function remains limited, particularly studies focused on African American women ^10^.

Blood pressure, hypertension, kidney function and CKD are complex, multifactorial traits influenced by a combination of genetic predisposition, molecular pathways, and environmental as well as social determinants ^11-21^. The analysis of mRNA expression provides insights into the integrated effects of these factors, reflecting both genetic regulation and cellular responses to environmental stimuli ^22-24^. Advances in high-throughput sequencing technologies have facilitated transcriptomic studies, enabling the identification of unique mRNA expression patterns that differentiate diseased from healthy individuals ^25-29^ and drug response ^30^.

However, these studies are predominantly conducted in European-ancestry populations ^31,32^, resulting in limited generalizability to other groups including African Americans ^33^. Furthermore, even within the African American population, women’s health remains understudied ^34^, despite the unique hormonal, biological, and environmental factors that may influence the pathogenesis of hypertension and its downstream effects on kidney function.

Kidney function, often measured as estimated glomerular filtration rate (eGFR), is a well-established marker of renal health ^35,36^. Hypertension is known to affect eGFR ^1^, but the biological mechanisms linking hypertension, kidney function, and genetic variation remain poorly understood.

In this study, we aim to address these gaps by leveraging transcriptomic data to investigate the intersection of hypertension and kidney function in African American women. Specifically, the project seeks to define a shared transcriptomic profile associated with hypertension and kidney function. By integrating transcriptomic and genomic data, this project will further our understanding of the molecular mechanisms linking hypertension, blood pressure and kidney function in African American women. This approach not only addresses important knowledge gaps in the genomics of hypertension and kidney health but also provides a robust foundation for experimental work that could yield mechanistic insights. These insights would inform precision medicine strategies and contribute to efforts aimed at reducing health disparities in this underserved population.

## MATERIAL AND METHODS

The data utilized in this study were derived from the GENomics, Environmental FactORs, and Social DEterminants of Cardiovascular Disease in African Americans Study (GENE-FORECAST) ^37-39^. The study was approved by the Institutional Review Board of the National Institutes of Health and conducted in compliance with relevant local laws and institutional guidelines. All participants provided written informed consent prior to their inclusion in the study.

### Phenotype Data

GENE-FORECAST is a comprehensive research platform designed to utilize a multi-omics systems biology approach for detailed analysis of minority health and disease in African Americans. It established a cohort of self-identified U.S.-born African American 344 females and 147 males, aged 21-65 from the Washington D.C. area. The 344 females were randomly divided into two equal groups, each consisting of 172 samples, to create discovery and validation datasets. This approach allowed for the analyses to be conducted in the discovery dataset and subsequently validated in the validation dataset; the characteristics of the individuals are outlined in Table 1. Hypertension (HTN) status was assessed by averaging three blood pressure (BP) readings and considering the use of antihypertensive medications. Participants were classified as hypertensive if their systolic blood pressure (SBP) was ≥140 mmHg, their diastolic blood pressure (DBP) was ≥90 mmHg, or if they were on medication for high BP. Individuals were categorized as normotensive if they had optimal BP levels (SBP ≤120 mmHg and DBP ≤80 mmHg) without requiring BP-lowering medications. Estimated glomerular filtration rate (eGFR) was calculated using the creatinine-based equation developed by the Chronic Kidney Disease-Epidemiology Collaborative Group (CKD-EPI), adhering to guidelines from the National Kidney Foundation and the American Society of Nephrology Task Force ^40^. The race-neutral CKD-EPI equation was used; this equation demonstrated an improved correlation between measured and calculated GFR ^41^. The eGFR values ranged from 48 to 145 mL/min/1.73 m^2^, with nine participants registering values below 60.

**Table 1:**
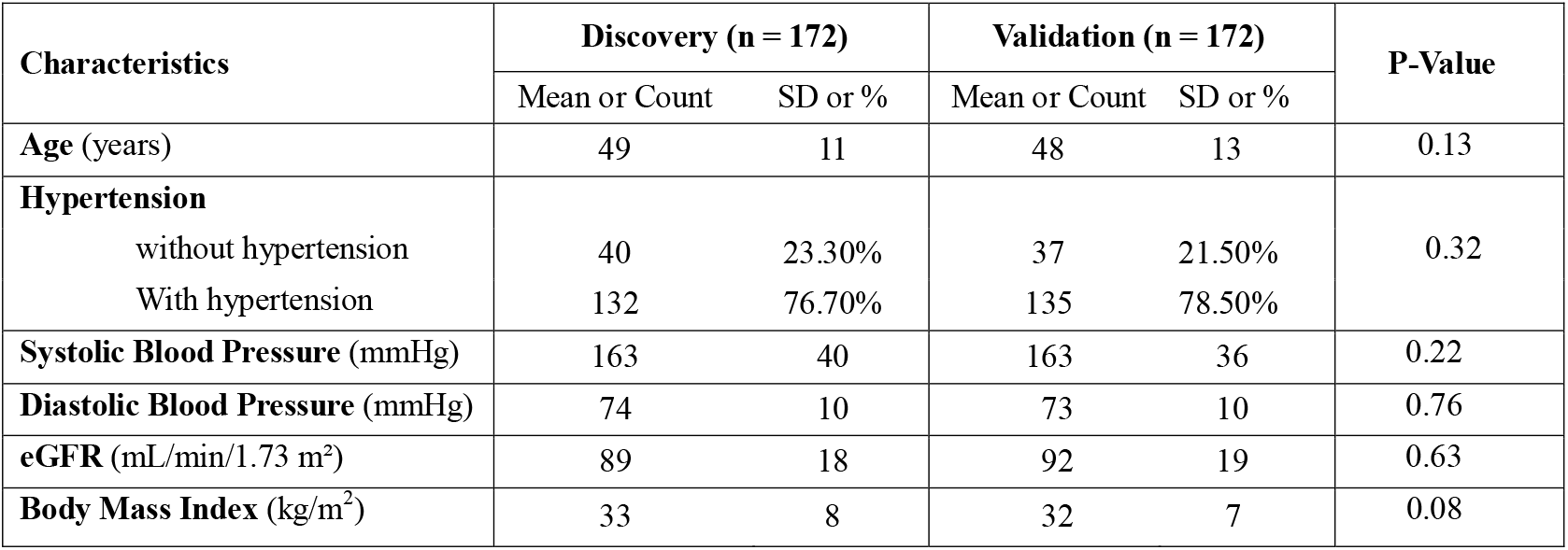
Characteristics of the GENE-FORECAST female subjects in the discovery and validation datasets.

| Characteristics | Discovery (n = 172) |  | Validation (n = 172) |  | P-Value |
| --- | --- | --- | --- | --- | --- |
|  | Mean or Count | SD or % | Mean or Count | SD or % |  |
| Age (years) | 49 | 11 | 48 | 13 | 0.13 |
| **Hypertension** |  |  |  |  |  |
| without hypertension | 40 | 23.30% | 37 | 21.50% | 0.32 |
| With hypertension | 132 | 76.70% | 135 | 78.50% |  |
| **Systolic Blood Pressure** (mmHg) | 163 | 40 | 163 | 36 | 0.22 |
| **Diastolic Blood Pressure** (mmHg) | 74 | 10 | 73 | 10 | 0.76 |
| **eGFR** (mL/min/1.73 m<sup>2</sup>) | 89 | 18 | 92 | 19 | 0.63 |
| **Body Mass Index** (kg/m<sup>2</sup>) | 33 | 8 | 32 | 7 | 0.08 |

The characteristics of the 147 males are outlined in **Table 2**. The male and female groups were similar in age, with both having a mean age of 49 years. Hypertension prevalence was high in both groups but slightly higher in males (77.60%) compared to females (76.70%). Both groups exhibited similar mean SBP (163 for both sexes) and mean of DBP (74 for both sexes). Regarding kidney function, males had a slightly higher mean eGFR value (91) compared to females (89). BMI was marginally lower in males (mean: 32) compared to females (mean: 33). In summary, the male and female groups showed comparable characteristics.

**Table 2:**
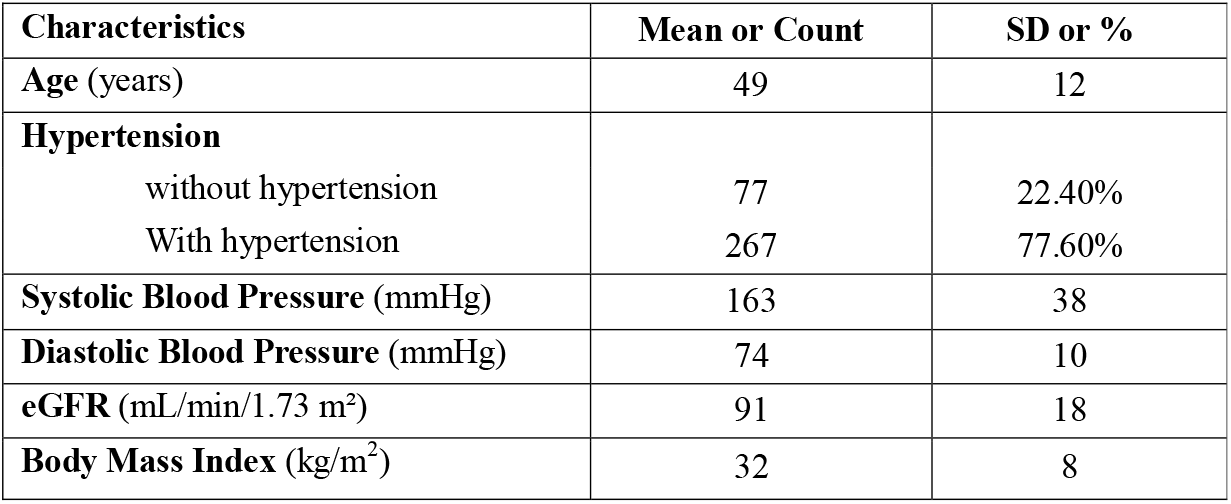
Characteristics of the 147 GENE-FORECAST male subjects.

| Characteristics | Mean or Count | SD or % |
| --- | --- | --- |
| Age (years) | 49 | 12 |
| **Hypertension** |  |  |
| without hypertension | 77 | 22.40% |
| With hypertension | 267 | 77.60% |
| **Systolic Blood Pressure** (mmHg) | 163 | 38 |
| **Diastolic Blood Pressure** (mmHg) | 74 | 10 |
| **eGFR** (mL/min/1.73 m<sup>2</sup>) | 91 | 18 |
| **Body Mass Index** (kg/m<sup>2</sup>) | 32 | 8 |

The male subset (n=147) was not divided into discovery and validation subsets due to the smaller sample size, which would have reduced statistical power and hindered meaningful interpretation. Instead, male data were used to provide a complementary perspective and to contrast differentially expressed mRNA identified in males with those in females. This strategy ensures the male data serve as a reference to isolate female-specific transcriptomic profiles while maximizing the utility of the available data without compromising its interpretability.

### Transcriptome Data

The transcriptome dataset comprises messenger RNA sequencing from whole blood samples. Total RNA was isolated using the MagMAX™ RNA Isolation Kit for Stabilized Blood Tubes, following the manufacturer’s instructions (Life Technologies, Carlsbad, CA). Indexed cDNA sequencing libraries were prepared from total RNA using Illumina’s TruSeq kits, with ribosomal RNA (rRNA) depletion performed during the process.

Samples were sequenced using paired-end reads on the Illumina HiSeq2500 and HiSeq4000 platforms, achieving a minimum depth of 50 million reads per sample. The quantification of mRNA expression was carried out using the Genotype-Tissue Expression (GTEx) bioinformatics pipeline, developed by the Broad Institute, with detailed documentation available on GitHub ^42^. The pipeline utilized tools such as FastQC (v0.11.5), STAR (v2.4.2a), samtools (v1.3), bamtools (v2.4.0), picard-tools (v2.5.0), and RSEM (v1.2.22). Transcripts with expression levels below 2 counts per million (CPM) in fewer than three samples were excluded.

Normalization of the expression data was performed using the Trimmed Mean of M-values (TMM) method ^43^, which is particularly suited for read count data. Principal component analysis (PCA) was employed to identify and remove outlier samples and transcripts. After applying these quality control measures, a total of 17,947 protein-coding mRNAs were retained for downstream statistical analyses.

## STATISTICAL ANALYSES

### Differential Expression Analyses in females

The R library *edgeR* was utilized to identify mRNAs differentially expressed between hypertensives and normotensives (step 1 in Figure 1) and between the upper and lower tertiles of eGFR (step 2 in Figure 1). All 17,947 protein-coding mRNAs that passed quality control filtering were considered for these analyses. *edgeR* fits a negative binomial model to the mRNA read counts (i.e. expression) and computes likelihood ratio tests for the coefficients in the model. The model was adjusted for age in both the discovery and validation datasets. mRNAs were considered significantly differentially expressed (DE) if the Benjamini-Hochberg (BH) ^44^ false discovery rate (FDR) adjusted p-value ≤ 0.05 in the discovery dataset, the nominal p-value ≤ 0.05 in the validation dataset and the log fold change (logFC) of the difference is in the same direction, in both datasets.

**Figure 1:**
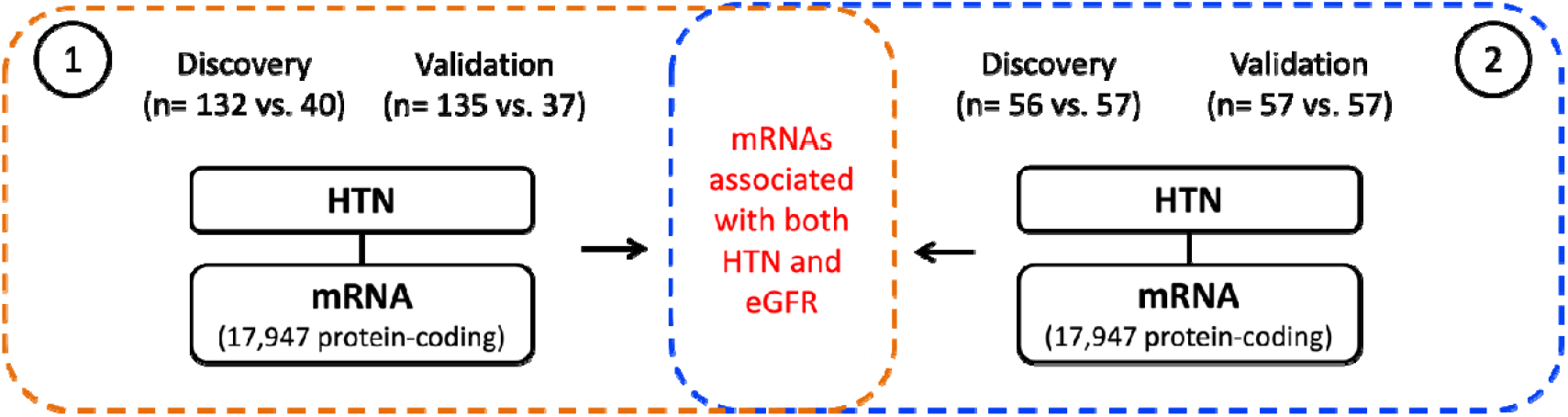
Overview of the analytical workflow and outcome in the female group: (1) Identification and validation of mRNAs that are differentially expressed by HTN status (hypertensives vs. normotensives) (2) Identification and validation of mRNAs differentially expressed between the lower and upper tertiles of eGFR. These analyses aim to uncover shared transcriptomic profiles between hypertension and kidney function.

The relationship between HTN status and mRNA expression was evaluated in a set of 172 females (132 with HTN vs. 40 without HTN) and validated in a set of 172 females (135 with HTN vs. 37 without HTN).

For eGFR, the differential expression analysis targeted the lower and upper tertiles of its distribution. This approach was chosen because the extremes of the distribution are likely to reveal mRNAs and pathways with strong and biologically significant associations. Such associations may be more pronounced compared to those observed across the entire range of eGFR. Moreover, focusing on these extremes enhances statistical power, as distinct differences are more readily detectable in extreme groups. This strategy helps to reduce noise and variability, which could otherwise obscure meaningful findings in a heterogeneous study population. The association between eGFR and mRNA expression was evaluated in a set of 113 females (57 in the lower tertile vs. 56 in the upper tertile) and validated in a set of 114 females (135 with HTN vs. 37 without HTN).

### Comparative Analysis of Male and Female Transcriptome Profiles Associated with Hypertension and eGFR

The analyses performed in the female cohort were similarly conducted in the male group, enabling a comparative evaluation of overlapping and distinct mRNA transcripts associated with hypertension and eGFR between the sexes.

First, the relationship between HTN status and mRNA expression was evaluated in a set of 147 males (114 with HTN vs. 33 without HTN). Then, the association between eGFR and mRNA expression was evaluated in a set of 97 males (48 in the lower tertile vs. 49 in the upper tertile). In both analyses, the negative binomial model was adjusted for age; mRNAs were considered significantly differentially expressed if the FDR adjusted p-value ≤ 0.05.

Finally, the results in the males were contrasted to those of females to identify overlap and female specific transcriptome profiles.

### Pathway Enrichment Analysis

Pathway enrichment analysis was conducted to identify biological pathways significantly represented within the list of mRNAs shared between HTN and eGFR. This analysis is important for uncovering the underlying molecular mechanisms that link blood pressure regulation and kidney function. By mapping these mRNAs to known pathways, the enrichment analysis provides insights into the interconnected biological processes related to both vascular and kidney function. This analysis was conducted using a function of the R library *edgeR* which runs hypergeometric tests, with mRNAs from our results mapped to known genes cataloged in the Kyoto Encyclopedia of Genes and Genomes (KEGG) database ^45^.

## RESULTS

### Genes associated with hypertension and eGFR in females and males

The mRNA differential expression analysis identified 3,320 mRNAs with statistically significant differential expression between hypertensive and normotensive females in the discovery dataset (FDR-adjusted p-value ≤ 0.05). Of these, 1,137 mRNAs were successfully validated in the validation dataset (p-value ≤ 0.05). A detailed list of these differentially expressed mRNAs, including their logFC values and corresponding p-values, is provided in **Supplemental Table T1A**.

The eGFR analysis revealed 3,320 mRNAs significantly differentially expressed between the lower and upper tertiles of eGFR in the female discovery dataset. Among these, 469 mRNAs were validated in the validation dataset. The comprehensive list of these eGFR-associated mRNAs, along with their respective logFC values and p-values, is available in **Supplemental Table T1B**.

**Figure 2A** illustrates the overlap of 400 mRNAs that were common between hypertension-associated (1,137) and eGFR-associated (469) mRNAs. The 400 mRNAs associated with both hypertension and eGFR, in the female set, are reported in **Supplemental Table T1C**.

**Figure 2:**
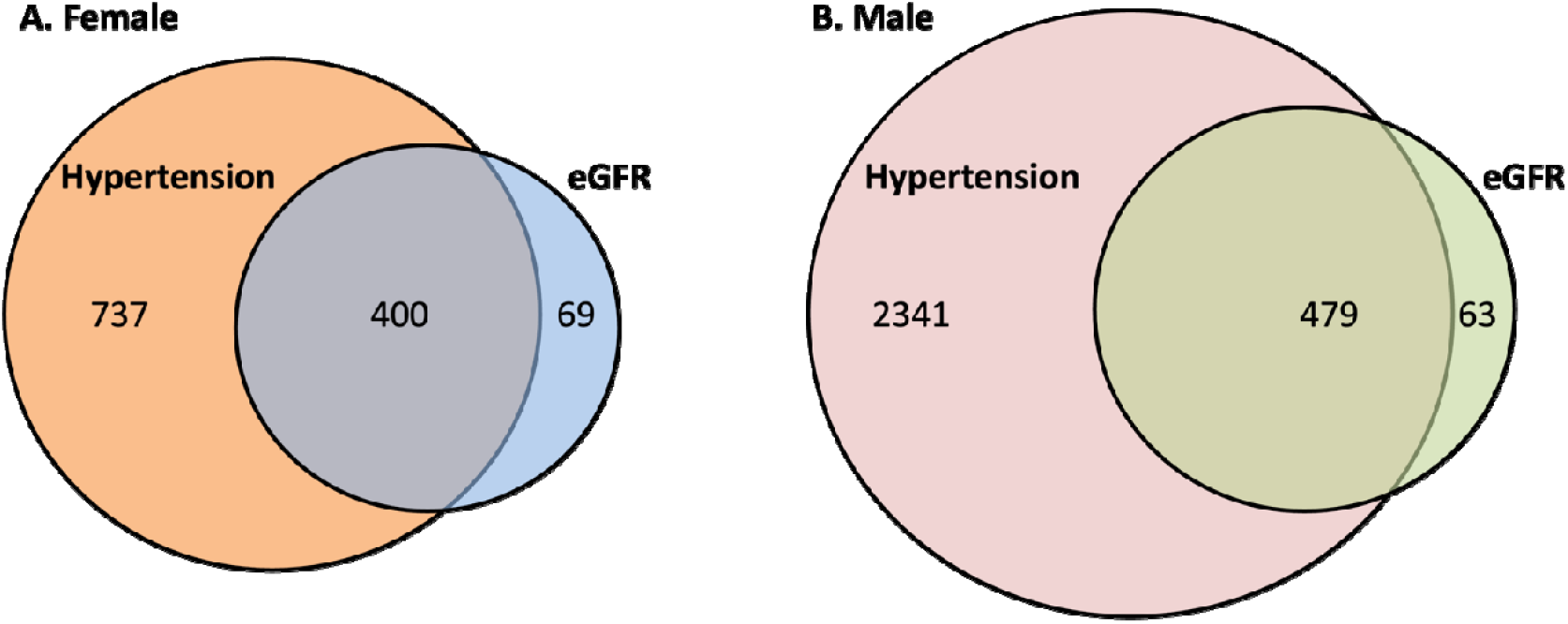
Differentially expressed mRNAs common to HTN and eGFR in females (A) and males (B).

Similar analyses, in the male group (n = 147) identified 2,820 mRNAs differentially expressed (FDR p-value ≤ 0.05) between hypertensive and normotensive subjects. A total of 542 mRNAs were differentially expressed between lower and upper tertile of eGFR. The comprehensive list of mRNAs associated with hypertension and eGFR are **Supplemental Table T1D** and **T1E**, respectively.

**Figure 2B** illustrates the overlap of 479 mRNAs that were common between hypertension-associated (2,820) and eGFR-associated (542) mRNAs. The 479 mRNAs associated with both hypertension and eGFR, in the male set, are reported in **Supplemental Table T1F**.

### Sex-specific comparisons of mRNA expression profiles associated with hypertension and eGFR

To identify female-specific transcriptomic profiles, the lists of mRNAs associated with hypertension and eGFR in females were compared to those identified in males. The results of these comparisons are reported graphically in **Figure 3**. A total of 1,137 mRNAs were associated with hypertension in females, with 89 unique to females and 1,048 overlapping with males. For eGFR, 689 mRNAs were identified in females, with 148 specific to females and 321 shared with males. The final comparison revealed 400 mRNAs associated with both hypertension and eGFR in females, with 95 unique to females and 305 shared with males.

**Figure 3:**
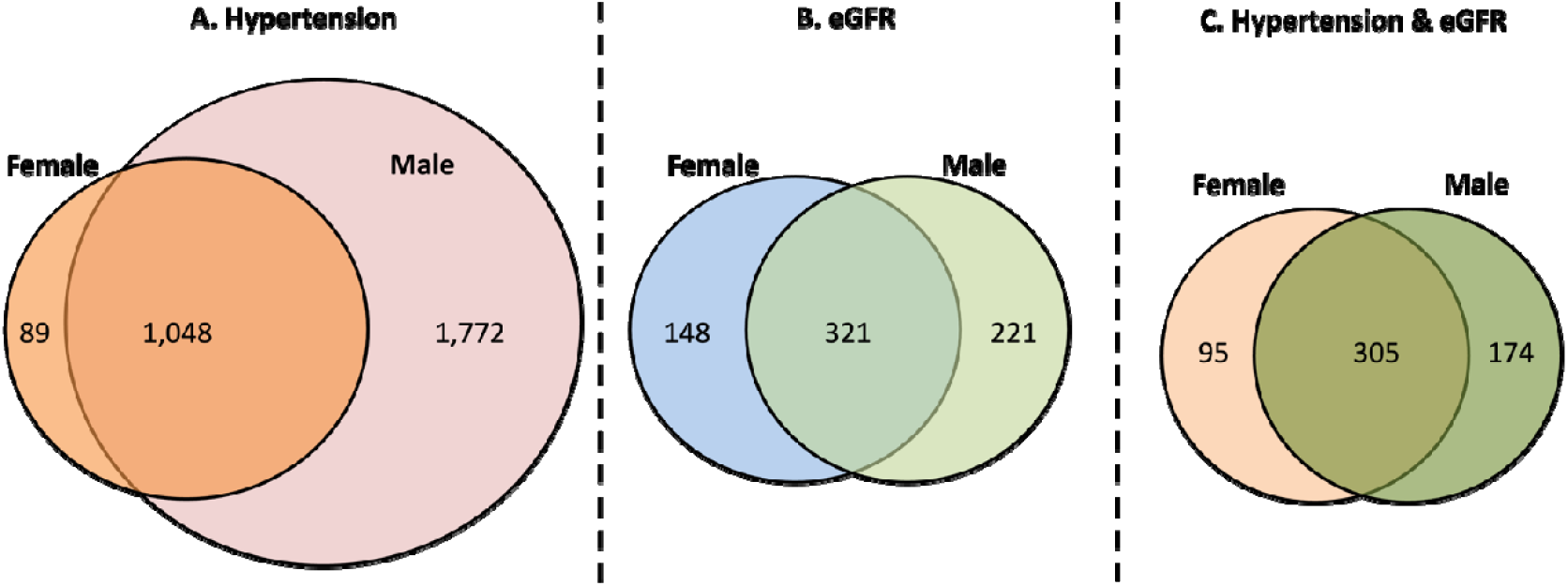
Venn diagrams summarizing the comparison of mRNAs associated with hypertension and eGFR between females and males. Panel A shows mRNAs associated with hypertension, with 89 unique to females. Panel B represents mRNAs associated with eGFR, with 148 unique to females. Panel C displays mRNAs associated with both hypertension and eGFR, with 95 unique to females.

### Female-specific mRNAs associated with both hypertension and eGFR

A total of 95 were associated with hypertension and eGFR in females only (Figure 4).

**Figure 4:**
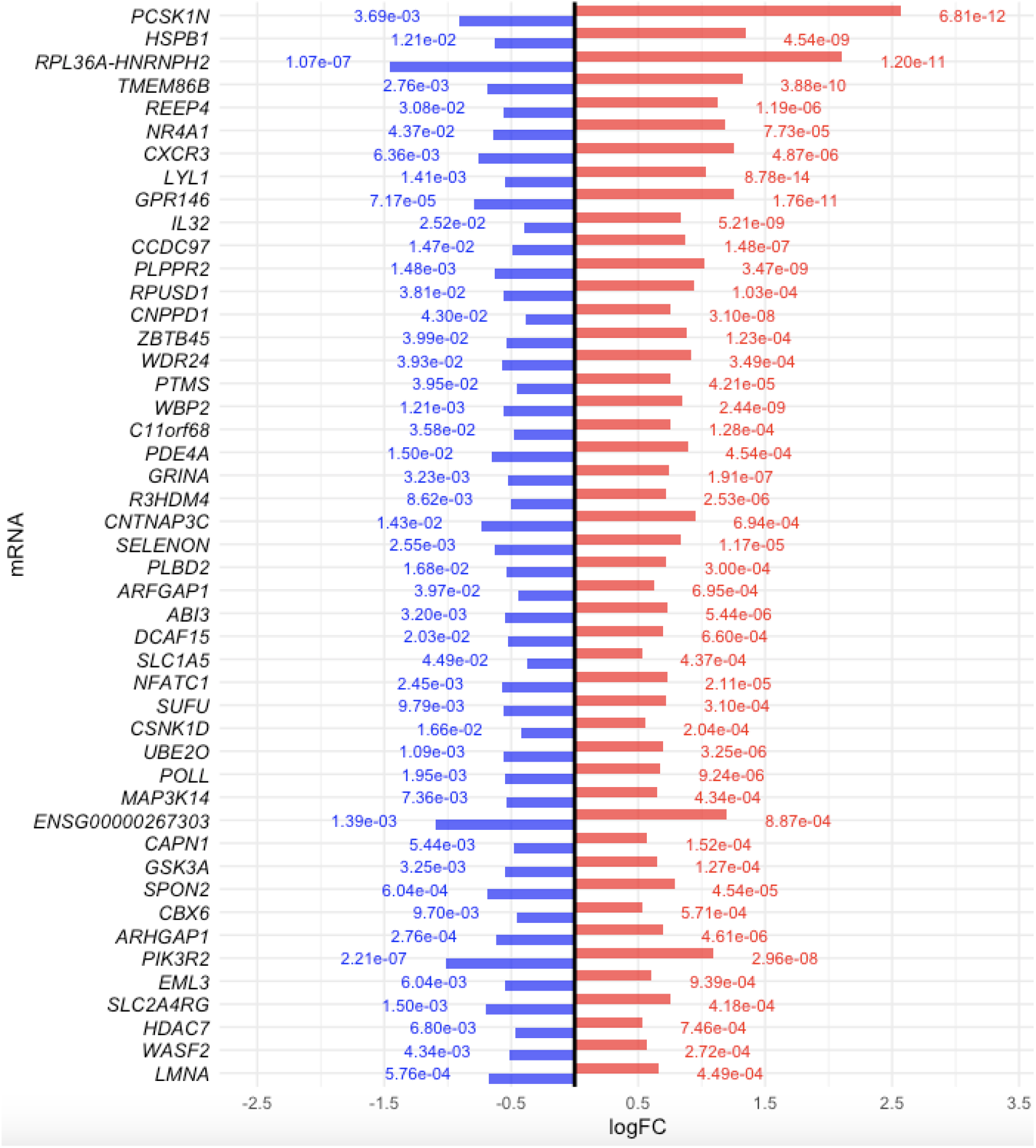
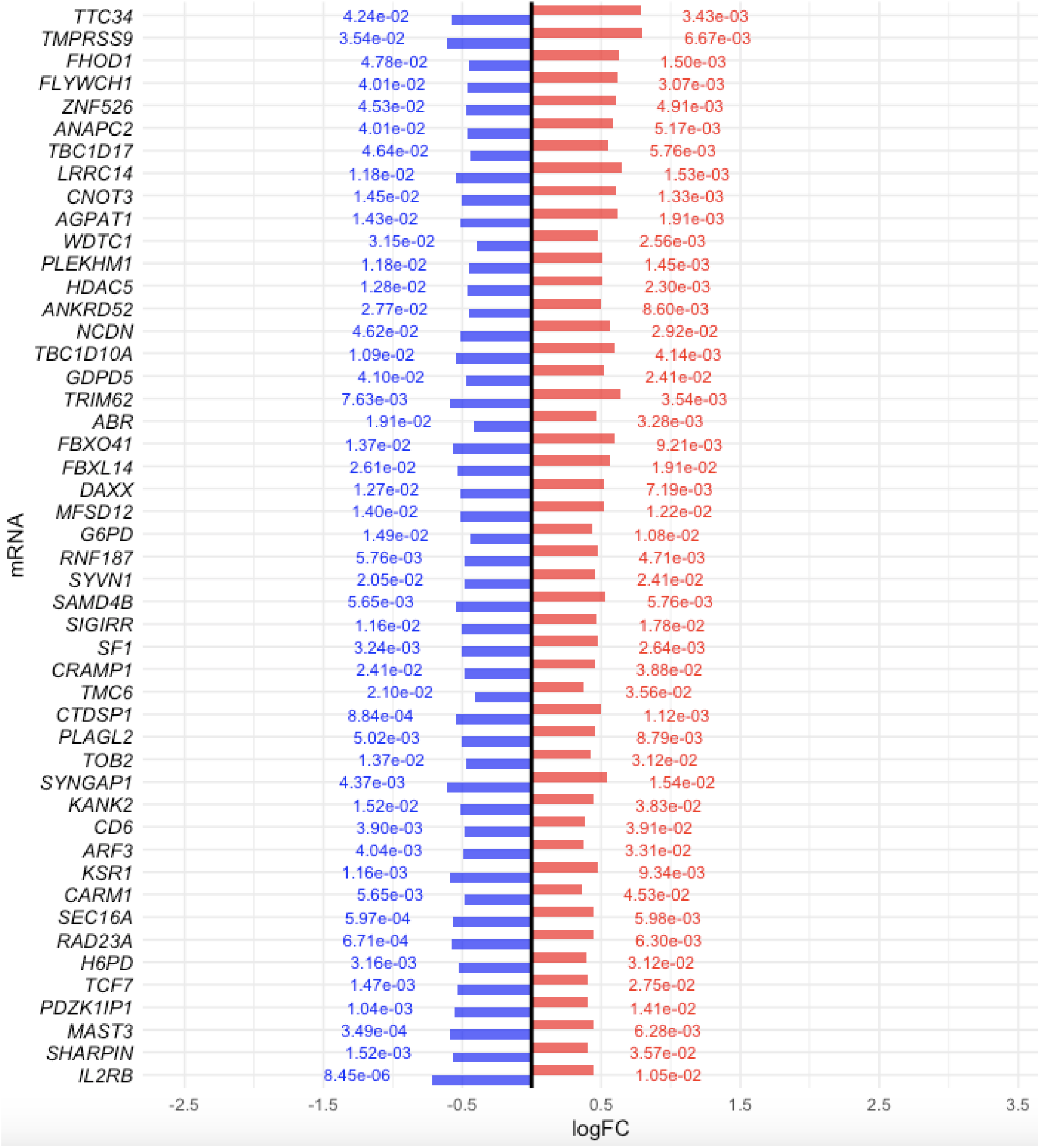
Differential expression analysis of the 95 female-specific mRNAs associated with both hypertension and eGFR. The horizontal bars represent the log2 fold changes (logFC) for each mRNA, colored by trait (red for hypertension and blue for eGFR). The values next to the bars indicate the corresponding FDR-adjusted p-values.

### Pathways enriched in the female-specific set of mRNAs associated with both hypertension and eGFR

Pathway enrichment analysis, summarized in Figure 5 which reports pathways with p-value ≤ 0.05, revealed significant involvement of multiple biological pathways, including apoptosis, immune signaling (e.g., C-type lectin receptor signaling), and metabolic processes (e.g., central carbon metabolism). These findings highlight key mechanisms potentially underlying the associations between transcriptomic profiles, hypertension, and kidney function.

**Figure 5:**
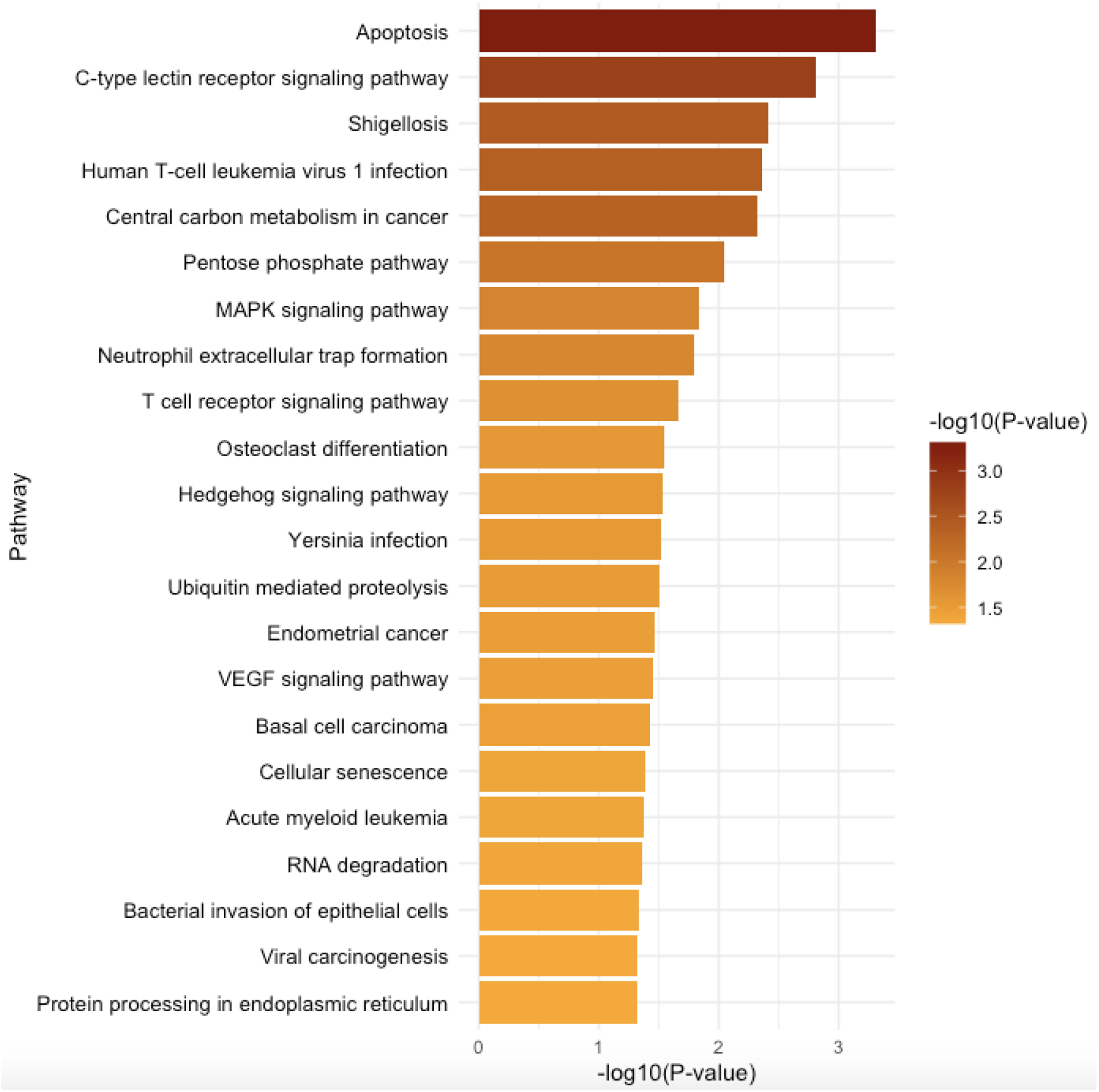
Pathways enriched in the set of 95 mRNAs associated with hypertension and kidney function. The pathways are ranked by p-value. The color gradient represents p-values, ranging from the lowest to the highest on a -log10 scale.

## DISCUSSION

### Summary

The primary focus of this study is to identify female-specific whole-blood transcriptomic profiles associated with hypertension and kidney function, measured as eGFR, in African Americans. To achieve this, genes associated with hypertension and eGFR in 344 females were compared to those from 147 males. The analyses identified 95 female-specific genes that were upregulated in women with hypertension and downregulated in women in the lower tertile of eGFR. These findings highlight genes such as TGF-β1 and their involvement in signaling pathways related to fibrosis, inflammation, and endothelial dysfunction. Additionally, the results underscore the role of genes like PNPLA2 in altered lipid metabolism, which exacerbates oxidative stress and kidney damage. Further relevant discoveries include immune pathways involving genes such as IL32 and TNFα, which amplify inflammation and contribute to renal injury.

### Genes and Signaling Pathways Driving eGFR Decline

#### TGF-β1 and eGFR Signaling

The interaction between TGF-β1 signaling and declining eGFR highlights their contribution to fibrotic and inflammatory responses in CKD ^46^. Increased TGF-β1 expression is associated with fibroblast activation, epithelial proliferation, and extracellular matrix (ECM) deposition through upregulation of SMAD3 phosphorylation, a key downstream mediator of TGF-β1-induced fibrosis ^47,48^. In our study, African American women with hypertension demonstrated upregulation of TGFB1, which was associated with lower eGFR, consistent with the profibrotic role of TGF-β1 observed in previous studies ^49^. Additionally, elevated expression of inflammatory markers was correlated with eGFR decline, perpetuating interstitial inflammation ^50-52^.

#### Cytoskeletal Dynamics and Rho GTPases

Cytoskeletal remodeling plays a significant role in maintaining the integrity of glomerular podocytes, mesangial cells, and tubular epithelial cells ^53^. Rho GTPases regulate cytoskeletal stability, but their dysregulation specifically RhoA and Rac1 leads to podocyte effacement, glomerular sclerosis, and impaired kidney filtration ^54^. Our study observed a downregulation of ARHGDIA (Rho GDP dissociation inhibitor alpha), a key regulator of Rho GTPase activity, consistent with evidence that disrupted cytoskeletal dynamics exacerbate endothelial permeability and fibrosis ^55^.

### Metabolic and Immune Dysregulation

#### Lipid and ATP Metabolism

The regulation of lipid metabolism through PNPLA2 (adipose triglyceride lipase, ATGL) adds another layer of complexity, as lipid homeostasis is essential to prevent CKD-related complications ^56^. In our results, PNPLA2 downregulation was significantly associated with lower eGFR, consistent with its role in hydrolyzing triglycerides to prevent ectopic lipid accumulation ^57^. Dysregulated lipid metabolism promotes the accumulation of reactive oxygen species (ROS), which exacerbates podocyte injury through cytoskeletal disruption ^58^. The role of lipid metabolism in CKD progression is further influenced by adipokines, such as leptin and adiponectin ^59^. Elevated leptin levels promote endothelial dysfunction and oxidative stress, while reduced adiponectin levels exacerbate inflammation and insulin resistance ^60^.

Similarly, adenosine triphosphate (ATP) metabolism plays an important role in regulating kidney function through its role in tubuloglomerular feedback (TGF) ^61^. Adenosine, a breakdown product of ATP, binds to adenosine receptors (A1AR and A2AR) to regulate afferent arteriolar vasoconstriction and medullary vasodilation ^62^. Our results suggest that chronic adenosine signaling contributes to eGFR decline, likely by enhancing sodium reabsorption and promoting inflammation. The observed downregulation of ATP6V0C (ATPase H+ transporting V0 subunit C) in females on the lower tertile of eGFR, along with its upregulation in hypertensives individuals, is consistent with impaired ion transport and acid-base regulation.

#### Immune-Driven Mechanisms of Hypertension and CKD Progression

The study revealed a set of genes including IL32, IL2RB, CD7, and TNFSF12 known to contribute to inflammation, fibrosis, and kidney dysfunction in hypertension and CKD ^63-65^. These genes, upregulated in hypertensives and downregulated in females on the lower eGFR tertile, are part of a connected network of immune regulation, shaping both adaptive and innate responses relate to disease.

At the center of this immune regulation, IL32 stands out for its capacity to amplify pro-inflammatory signals, particularly through the upregulation of IFN-γ, IL6, and TNFα, thereby inducing immune cell activation and recruitment ^66^. The observed elevated IL32 expression in hypertensive individuals in our analysis aligns with findings from studies linking it to kidney injury in diabetic kidney disease (DKD) and systemic inflammation in metabolic disorders. This highlights IL32’s role in bridging metabolic and immune pathways, creating a pro-inflammatory environment that primes renal tissue for further damage ^67^.

In parallel, IL2RB supports the expansion and persistence of T-cell responses, reinforcing the inflammatory cascade initiated by IL32 ^68^. Dysregulated IL2RB signaling enhances IFN-γ production, which drives the recruitment of pro-inflammatory CD8+ T cells and macrophages to renal tissues ^69^. This continuous influx of immune cells exacerbates endothelial damage, a process compounded by the involvement of CD7, a key regulator of immune cell trafficking ^70^. The upregulation of CD7 in hypertensives and downregulation in lower eGFR tertile, reflects its role in facilitating T-cell migration and amplifying cytokine signaling, thereby contributing to vascular inflammation and tubular apoptosis ^71,72^.

Further amplifying this immune dysregulation is TNFSF12 (encodes the protein TWEAK), which interacts with the Fn14 receptor to induce endothelial cell apoptosis and promote tissue remodeling ^73^. The activation of the TWEAK/Fn14 axis has been associated with glomerular injury and fibrosis ^74^. This is consistent with the negative correlation of the expression of TNFSF12 eGFR, observed in our analysis.

While IL32, IL2RB, CD7, and TNFSF12, are important players, it is important to recognize that the immune landscape in hypertension-related impaired kidney function CKD extends beyond these genes. Other immune-related genes and pathways, such as those involved in complement activation ^75^, chemokine signaling, and antigen presentation, may also play significant roles in disease progression ^76^. These additional players may interact with the identified genes to exacerbate immune responses, inflammatory signaling, or even provide compensatory anti-inflammatory mechanisms. This interconnected network underscores the importance of a multi-targeted therapeutic approach to modulate immune responses, reduce inflammation, and preserve kidney function in patients with hypertension.

### Strengths and limitations

This study leverages a robust transcriptomic analysis pipeline, utilizing high-throughput sequencing data and rigorous differential expression methodologies to identify mRNAs associated with hypertension and kidney function. The two-step validation approach in the female cohort ensures reproducibility and strengthens the reliability of findings. The comparative analysis between male and female cohorts provides valuable insights into sex-specific transcriptomic differences, and the integration of pathway enrichment analysis links observed mRNA profiles to relevant biological processes, offering mechanistic insights.

While the study’s strengths underscore its potential to provide meaningful insights into the transcriptomic landscape of hypertension and kidney function in African American women, several limitations warrant consideration to contextualize the findings and guide future research directions. Focusing on tails of the eGFR distribution, while increasing statistical power, may overlook subtler transcriptomic changes present in intermediate ranges. The Pathway enrichment analysis depends on existing pathway databases, which may not fully capture the nuances of less well-characterized or novel pathways relevant to hypertension and kidney function. Finally, the study design is cross-sectional, limiting the ability to infer causal relationships between identified mRNAs and the phenotypes of interest.

## Conclusions

This study provides critical insights into the transcriptomic landscape linking hypertension and kidney function in African American women. By identifying 95 female-specific genes associated with both traits, we highlight pathways involving TGF-β1, Rho GTPases, and PNPLA2 that underscore the roles of fibrosis, inflammation, lipid metabolism, and endothelial dysfunction in disease progression. Additionally, immune dysregulation through genes like IL32 and TNFα further connects hypertension and impaired kidney function, revealing potential therapeutic targets. These findings emphasize the importance of addressing sex-specific and population-specific molecular mechanisms to develop targeted interventions for reducing health disparities in chronic kidney disease. Future research should explore longitudinal designs and experimental validation to deepen our understanding of these complex interactions.

